# Transparency reduces predator detection in chemically protected clearwing butterflies

**DOI:** 10.1101/410241

**Authors:** Mónica Arias, Johanna Mappes, Charlotte Desbois, Swanne Gordon, Melanie McClure, Marianne Elias, Ossi Nokelainen, Doris Gomez

**Affiliations:** CEFE, Univ Montpellier, Univ Paul Valéry Montpellier 3, EPHE, IRD, Montpellier, France; Department of Biological and Environmental Science, Centre of Excellence in Biological Interactions, University of Jyvaskyla, P.O. Box 35, 40014 Jyvaskyla, Finland; Institut de Systématique, Evolution, Biodiversité, UMR 7205, CNRS, MNHN, Sorbonne Université, EPHE, 45 rue Buffon CP50, France; INSP, Sorbonne Université, CNRS, Paris, France

**Keywords:** aposematic, crypsis, detectability, Ithomiini, vision modelling, citizen science, bird, experiment

## Abstract

1. Predation is an important selective pressure and some prey have evolved warning colour signals advertising unpalatability (i.e. aposematism) as an antipredator strategy. Unexpectedly, some butterfly species from the unpalatable tribe Ithomiini possess transparent wings, an adaptation rare on land but common in water where it helps avoiding predator detection.

2. We tested if transparency of butterfly wings was associated with decreased detectability by predators, by comparing four butterfly species exhibiting different degrees of transparency, ranging from fully opaque to largely transparent. We tested our prediction using using both wild birds and humans in behavioural experiments. Vision modelling predicted detectability to be similar for these two predator types.

3. In concordance with predictions, more transparent species were almost never the first detected items and were detected less often than the opaque species by both birds and humans, suggesting that transparency enhances crypsis. However, humans could learn to better detect the most transparent species over time. Our study demonstrates for the first time that transparency on land likely decreases detectability by visual predators.

## Introduction

Predation is an important selective pressure and a strong evolutionary force shaping prey coloration. As a way to avoid predator detection, some prey have evolved colours and textures that mimic those of the background, hence rendering them cryptic (Endler, 1988). In midwater environments where there is nowhere to hide, crypsis can be achieved by different means, including transparency (Johnsen, 2014). Transparency is common in aquatic organisms where it has been shown to decrease detectability by visual predators, enabling prey to blend in with their environment (Kerfoot, 1982; Langsdale, 1993; Tsuda, Hiroaki, & Hirose, 1998; Zaret, 1972). By contrast, transparency is generally rare in terrestrial organisms, except for insect wings, which are made of chitin, a transparent material. The lack of pigments in these wings is sometimes accompanied by anti-reflective nanostructures that render them highly transparent, such as in cicadas and damselflies (Watson, Myhra, Cribb, & Watson, 2008; Yoshida, Motoyama, Kosaku, & Miyamoto, 1997). However, Lepidoptera (named after ancient Greek words for scale – *lepis* – and wing -*pteron*) are an exception as their wings are generally covered with colourful scales that are involved in intraspecific communication (Jiggins C. D., Estrada C., & Rodrigues A., 2004), thermoregulation (Miaoulis & Heilman, 1998), water repellence (Wanasekara & Chalivendra, 2011), flight enhancement (Davis, Chi, Bradley, & Altizer, 2012), and antipredator strategies such as crypsis (Stevens & Cuthill, 2006), masquerade (Suzuki, Tomita, & Sezutsu, 2014) and aposematism (i.e. advertisement of unpalatability, Mallet & Singer, 1987).

Ithomiini (Nymphalidae: Danainae), also known as clearwing butterflies, are some of the most abundant butterflies in Neotropical forests (Willmott, Willmott, Elias, & Jiggins, 2017). They are thought to be unpalatable due to the accumulation of pyrrolizine alkaloids collected from Asteraceae, Boraginaceae and Apocynaceae plants (Brown, 1984, 1985). Many of these clearwing butterflies represent classic examples of aposematic prey, whereby bright colour patterns – often with orange, yellow and black - advertise their unprofitability to predators (Mappes, Marples, & Endler, 2005; Poulton, 1890). Bright contrasting and aposematic coloration is likely to be the ancestral state in the group, since sister lineages (Tellerveni and Danaini) are opaque and aposematic (Freitas & Brown, 2004). However, transparency has evolved to some degree in approximately 80% of clearwing butterfly species, even though many retain minor opaque and colourful wing elements (Beccaloni, 1997; Elias, Gompert, Jiggins, & Willmott, 2008; Jiggins, Mallarino, Willmott, & Bermingham, 2006). Since transparency is often associated with crypsis, for example in aquatic organisms (Johnsen, 2014), transparency may have evolved in these butterflies to reduce their detectability by predators.

To determine if transparency in clearwing butterflies decreases detectability by visual predators, we compared predator detection of four Ithomiini species that differed in the amount of transparency of their wings (Fig S1): *Hypothyris ninonia* (largely opaque), *Ceratinia tutia* (transparent but brightly coloured), *Ithomia salapia* (transparent with a pale yellow tint) and *Brevioleria seba* (transparent without colouration other than a white band in the forewing). Given the proportion of light that is transmitted through the butterfly wing of the different species (Fig S2), we predicted that the opaque species *Hypothyris ninonia* should be easiest to detect, followed by the transparent but coloured *Ceratinia tutia.* Finally, the more transparent butterfly species *Ithomia salapia* and *Brevioleria seba* should be the least detectable. We tested our predictions using two complementary behavioural experiments, further supported by a vision modelling approach.

Detectability of butterflies was first tested using wild great tits (*Parus major*) as model bird predators. Great tits are highly sensitive to UV wavelengths (UVS vision in Ödeen, Håstad, & Alström, 2011). Their vision is similar to that of naturally occurring Ithomiini predators like the houtouc motmot (*Momotus momota*, Pinheiro, Medri, & Salcedo, 2008), the fawn-breasted tanager (*Pipraeidea melanonota*, (Brown & Neto, 1976) or the rufous-tailed tanager (*Ramphocelus carbo* (Brower, Brower, & Collins, 1963). However, great tits are naïve to ithomiine butterflies and they do not associate their colour patterns to toxicity. As bird responses are the result of both prey detection and motivation to attack the prey, we performed behavioural experiments using human participants, which can prove to be useful in disentangling these two factors. Despite differences in colour perception between humans and birds (both of which are visual predators), humans have been found to be good predictors of prey survival in the wild (Penney, Hassall, Skevington, Abbott, & Sherratt, 2012). Finally, models of predator vision (both for birds and humans) were used to complement behavioural experiments and infer the relative detectability of each butterfly species based on their contrast against the background.

## Materials and Methods

### Butterflies used for the behavioural experiments

Specimens of the four Ithomiini species used in both experiments – which, in order of increasing transparency are *Hypothyris ninonia, Ceratinia tutia, Ithomia salapia aquina, Brevioleria seba* (see Figs S1, S2) *–* were collected in Peru in 2016 and 2017, along the Yurimaguas - Moyobamba road (-6.45°, -76.30°). Butterflies were kept dry in glassine envelopes until use. In behavioural experiments, a single real hindwing and a single real forewing were assembled into artificial butterflies using glue and a thin copper wire to attach the artificial butterfly to a substrate (see Fig S3 for an example). These artificial butterflies mimicked real Ithomiini butterflies at rest, with wings closed and sitting on plant leaves (a typical posture for resting butterflies).

### Behavioural experiments using wild birds

Behavioural experiments took place in August and September 2017 at the Konnevesi Research Station (Finland) under permit from the National Animal Experiment Board (ESAVI/9114/04.10.07/2014) and the Central Finland Regional Environment Centre (VARELY/294/2015). Thirty wild-caught great tits (*Parus major*) were used, including 3 juvenile and 10 adult females, and 8 juvenile and 9 adult males. Birds were caught using spring-traps and mist-nets, individually marked with a leg band and used only once. Each bird was kept individually in an indoor cage (65x65x80 cm), with a 12:12 photoperiod. Birds were fed with peanuts, sunflower seeds, oat flakes and water *ad libitum*, except during training and experiments. During training, birds were given mealworms (see Training section). Birds were deprived of food for up to 2 hours before the experiment to increase their motivation. Most birds were kept in captivity for less than a week, after which they were released at their capture site.

#### Training

In their indoor cage, birds were taught that all four species of butterflies were similarly palatable by offering them wings of four butterflies (one of each species) with a mealworm attached to the copper wire. Butterfly wings used for training were laminated with transparent thin plastic to minimize damage so that these wings could be re-used when possible. As birds typically do not consume butterfly wings but only butterfly body (here dead mealworm), wing toxicity or unpalatability did not influence their training. Butterflies were presented to the birds in the absence of any vegetation during training. When birds had eaten all four prey items (one of each species), a new set was presented. Training ended when birds had eaten 3 sets of butterflies. No time constraint was imposed for training and most birds completed it in less than 4 hours.

In order to familiarise birds with the experimental set-up, which was novel to them, they were released in the experimental cage by groups of two to four birds for approximately one hour the day before the experiment. Oat flakes, seeds and mealworms were dispersed over leaves and vegetation to encourage searching for edible items in locations similar to where butterflies would be placed during the experiment.

#### Experiments

The experimental set-up consisted of a 10m x 10m cage that had tarpaulin walls and a ceiling of whitish dense net that let in natural sunlight. Butterflies were disposed in a 5 x 5 grid, delimited by poles all around the borders and a rope defining rows and columns (see Fig S4). Two extra poles were placed in the grid centre to increase the appeal of this area for birds. Five specimens of each species (20 specimens in total) were placed in the grid, one per cell, following a block randomization for each row and column and ensuring that all species were evenly represented along the grid. This random configuration was reshuffled between trials (i.e. randomized block design). The 5 cells closest to the observer were left empty as birds tended to avoid this area. For each trial, an observer, hidden to the birds, watched from outside the cage through a small window and took notes of which butterfly species were attacked and in which order. A GoPro camera also recorded the experiments. A butterfly was considered detected only if a bird directly approached it to grab, including when the attack failed. No bird was seen hesitating the attack once it started it. Experiments took place between 9 am and 5 pm. Before each trial, the radiance of ambient light (coming from the sun and sky) was measured using an Ocean Optics spectrophotometer in the same location each time. We computed the total radiance (TR) over the range of 300-700 nm of bird spectral sensitivity to account in statistical analyses for the level of ambient light intensity associated to each experimental trial. Further information on weather conditions (cloudy, sunny, etc) were also noted. Experiments ended when a bird had eaten half of the available butterflies (ie. 10 butterflies) or after 2 hours, whichever happened first. Wings were occasionally re-used if they had not been damaged.

After the experiment, the probability of a bird being present in a given grid cell was calculated as a proxy for the probability of an attack occurring in that grid cell. To do so, 10-minute intervals from each recording, selected based either on the maximum attack rate or when the bird was seen actively exploring the cage, were revised so as to calculate the proportion of time birds spent on poles situated next to each cell. A total of 87% of all attacks started from the pole closest to the grid cell, while all other attacks were initiated from a pole situated only one grid cell further away. Thus, the probability of visiting a given cell was the sum of the time spent on each neighbouring pole, divided by the number of either “close” (immediately next to) or “distant” (one grid cell removed) neighbouring cells and multiplied by either 0.87 or 0.13 (depending on the distance to the pole). As such, those cells closest to the poles and those at the edge of the cage were most likely to be visited by birds.

#### Statistical analyses

Differences in the total number of butterflies of each species that were attacked were compared by fitting generalised linear mixed effect models (GLMM), with bird identity as a random factor. A binomial distribution was used for the response variable (attacked or not), and the butterfly species, butterfly size, the probability of being attacked for a given cell, trial duration, age and sex of the bird, time to first attack, first butterfly species found, weather (as a qualitative variable), and total radiance (TR) were all selected as explanatory variables. The best fitting model was selected based on minimization of Akaike’s Information Criteria (AIC), assuming that models differing by two units or less were statistically indistinguishable (Anderson, Burnham, & White, 1998). Coefficients and standard errors were computed using a restricted maximum likelihood approach and a Wald z test was used to test for factor significance.

Most birds fed willingly on all butterflies located on the borders of the grid. Given that butterfly species distribution was random and reshuffled between trials, the four species were similarly represented in those cells (Fig S5), so no bias was expected. The order of attack for each species was tested by ranking the “inconspicuousness” of each butterfly species based on the order in which butterflies were found and how many of them were detected (Ihalainen, Rowland, Speed, Ruxton, & Mappes, 2012). To do so, the position of each butterfly that was attacked for each species and the total number of non-attacked butterflies of each species multiplied by 11 (i.e. the maximum number of butterflies that could be found + 1) were added. This inconspicuousness rank distinguishes between those species found first and in higher numbers (lower values of inconspicuousness) from those found last and in lower numbers (higher values of inconspicuousness). For example, if a bird captured two *H. ninonia* second and fifth in the sequence of captured prey, this species gets a rank value of 2+5+3x11=40 for that trial. We fitted a linear mixed effect model to test for differences in rank for each species, assuming a normal distribution, with rank as the response variable, bird individual as a random factor and butterfly species, age and sex of the bird, date, time until first attack, first butterfly species found, weather as a qualitative variable, and total radiance (TR) as explanatory variables. Again, the best fitting model was selected using AIC minimization.

We also tested whether differences in the rank of species inconspicuousness could be due to a differential attack probability for each species, i.e. whether species more likely to be attacked were more often placed on cells more likely to be visited. To do so, the probability of a bird being in proximity to a grid cell containing one of the five artificial butterflies of each species was averaged for each trial. An ANOVA was then used to compare the probability of attack for each butterfly species.

Finally, to test whether birds created a “search image” (i.e. improved in finding butterflies of a given species), the number of butterflies of each species found consecutively was counted. Results were compared among butterfly species using a χ^*^2^*^ test. Additionally, whether finding some species improved the bird’s ability to find others was tested. For each combination of two species, we calculated how many times a butterfly of species 1 was found after a butterfly of species 2. Differences between them were tested using a χ^*^2^*^ test. All analyses were performed in R (R Foundation for Statistical Computing, 2014).

### Behavioural experiments using human participants

Between mid-November and early December 2017, visitors of the Montpellier botanical garden (France) were invited to take part in an experiment where they searched for artificial butterflies. Before each trial, participants were shown pictures of various ithomiine butterflies, both transparent and opaque, but of species different than those used in the experiments, so as to familiarize them with what they would be searching for. Anonymous personal data was collected from each participant, including gender, age group (A1: <10 years, A2: 11-20 y, A3: 21-30 y, A4: 31-40 y, A5: 41-50 y and A6: >51 years), and vision problems. A participant did the experiment only once.

#### Experimental set-up

Artificial butterflies (N=10 of each of the four species, for a total of 40 butterflies) again consisted of one forewing and one hindwing assembled with copper wire and placed on leaves, but without the mealworm used in the bird experiments. These butterflies were set-up along two corridors in a forest-like understory habitat of similar vegetation and light conditions. Butterfly order followed a block randomisation, with five blocks each consisting of eight butterflies (i.e. two per species). This ensured that observers were similarly exposed to the four species all throughout the experimental transect. Whether a butterfly was placed on the left or right side of the corridor was also randomised. Both order and corridor side were changed daily. Participants could start the path from either end of the set-up and were given unlimited time to complete the trial. However, they could only move forward on the path. Only one participant was allowed in the path at any given time, and they were accompanied by an observer who recorded which butterflies were found. Trials ended when the participant had completed both corridors.

#### Statistical analyses

Differences in the total number of butterflies found for each species was tested by fitting GLMMs. A binomial distribution for the response variable (either found or not) was assumed, and participant identity was set as a random factor, butterfly species, first species found, butterfly position, corridor, left or right side of the path, time of day, gender and age of the participant, duration of the experiment, and their interactions, were all used as explanatory variables. A minimization of Akaike’s Information Criteria (AIC) was used to select the best model, assuming that models differing by two units or less were statistically indistinguishable (Anderson et al., 1998). Coefficients and standard errors were computed using a restricted maximum likelihood approach and a Wald z test was used to test for factor significance. Whether specific species were more frequently detected was also tested using a χ^*^2^*^ test. Similarly as for the experiments using birds, a GLMM was also fitted under the same conditions, but using the butterfly species inconspicuousness rank (similarly calculated as for the bird experiments) as a response variable.

Finally, whether humans found several butterflies of the same species consecutively (perhaps because they formed a “species search image”) was also tested. A χ^*^2^*^ test was used to compare the number of butterflies of each species that were found consecutively. Whether finding individuals of a species increased the likelihood of finding other species was also tested. For each combination of two species, we calculated how many times a butterfly of the first species was found immediately after a butterfly of a second species. Differences between the frequencies of these combinations were tested using a χ^*^2^*^ test, comparing observed results and the frequency at which each possible pair of species was placed consecutively in the original experimental setup. All analyses were performed in R (R Foundation for Statistical Computing, 2014).

Colour measurements and vision modelling can be found in electronic supplementary material (additional materials and methods).

## Results

### Behavioural experiments using wild birds

Birds took anywhere between 1 and 37 minutes (average: 7.54 ± 8.96 min) after release into the experimental cage before initiating an attack. For three of the birds, the experiment ended without having eaten 10 butterflies in the allocated 2 hours. The other 27 birds took between 11 and 112 minutes to attack 10 butterflies (mean time to attack 10 butterflies: 40.76 ± 26.23 min). Considering all trials, similar percentages of butterflies for each species was found by birds (54% of *H. ninonia* butterflies (the most colourful species), 48.7% for *C. tutia* (colourful but transparent species), 46.7% for *I. salapia* (yellow-tinted butterfly) and 49.3% of *B. seba* butterflies (most transparent species).

The model that best explained whether butterflies were attacked or not only included the probability of occupancy of a given grid cell by a bird, time of the first attacked and the grid cell occupied by the butterfly (AIC = 610.42, Delta AIC = 4.8142 with a model that additionally included butterfly species, Table S1). Butterflies were more likely to be attacked where birds visited most often (*z* = 2.93, *p* 0.003). By contrast, the inconspicuousness rank of a butterfly species was best explained by a model including average probability of occupancy by a bird and butterfly species as explanatory variables (AIC = 765.73, Delta AIC = 2.53 with a model including the species that was attacked first, Table S2). *H. ninonia*, which was the most colourful species, was usually detected in the first prey attacked (*t* = -3.15, Fig 1a, Table S2). Moreover, which species were found first closely matched their transmission properties: *H. ninonia,* followed by *C. tutia, I. salapia* and *B.seba* (*X*^2^ = 11.07, df = 3, *p* = 0.011, Table S3). When comparing species distribution along the grid, artificial butterflies that were attacked were in grid cells with moderate to high bird occupancy rate (*F* = 0.82, df = 3, *p* = 0.485, Fig S5).

**Figure 1.**
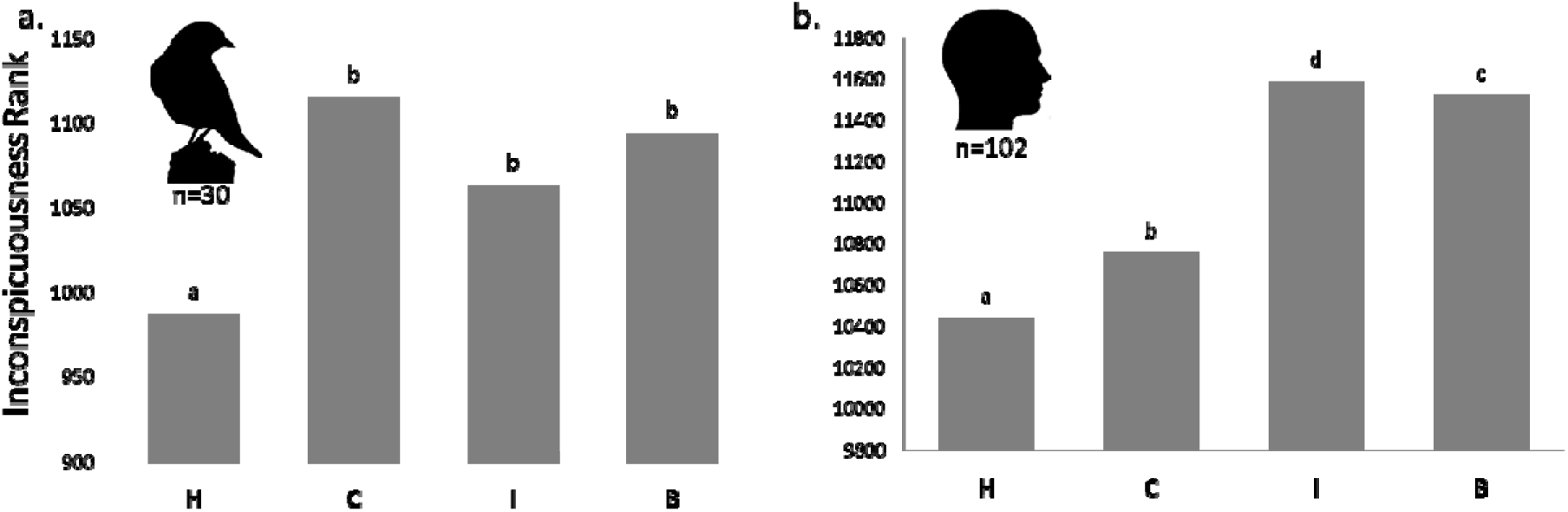
Sum of the inconspicuousness rank of each species for a) great tits and b) humans. Species for which butterflies were detected first and most often by birds or humans have lower values of “inconspicuousness rank”. Butterfly transparency increases from left to right: *H. ninonia* (H), *C. tutia* (C), *I. salapia* (I), and *B. seba* (B). Letters above the bars mean significant differences below 0.05.

Generally, birds did not find several butterflies of the same species consecutively (Fig S6a). In the rare instances that they did, no differences between species was found (*X*^*^2^*^ = 0.6, df = 3, *p* = 0.90) suggesting that birds did not form a “search image” for any of the butterfly species. No combination of species was attacked consecutively at high frequencies either (*X*^*^2^*^ = 10.88, df = 11, *p* = 0.45).

### Behavioural experiments using human participants

A total of 102 volunteers participated in the experiment (63 men and 39 women, with 10:11:21:18:31:11 in the A1:A2:A3:A4:A5:A6 age classes). 19 volunteers ran the experiment before 13h30, 35 between 13h30 and 16h, and 48 after 16h. Participants found between 5 and 28 of the 40 butterflies (12.75 ± 4.68 butterflies found per participant) and took between 7.5 and 37 minutes to walk both corridors (18.04 ±6.5 minutes spent in average per participant). For all trials combined, participants found 42.5% of *H. ninonia* butterflies (the most colourful species), 38% of *C. tutia* (colourful but transparent species), 23.54% of *I. salapia* (yellow-tinted butterfly) and 28.63% of *B. seba* butterflies (most transparent species).

Younger participants found more butterflies than older ones (number of butterflies: *z =* -2.34; butterfly species rank: *t =* -1.36). Additionally, participants found more butterflies earlier than later in the afternoon (number of butterflies: *z =* -2.80; inconspicuousness rank: *t = -* 1.77). However, this was most significant for younger participants (inconspicuousness rank: *t =* 1.32, Fig S7a). Generally the more time participants spent in the experiment, the higher the number of butterflies they found (number of butterflies: *z =* 5.21; inconspicuousness rank: *t =* 4.03), although this was most significant for women (number of butterflies: *z = -*2.96, inconspicuousness rank: -2.83, Fig S7b). There was a corridor effect, likely due to differences in the overall cover of vegetation (number of butterflies: *z =* 3.14; inconspicuousness rank: *t = -*3.52). Participants also found more butterflies at the end rather than at the start of the experiment (number of butterflies*: z =* 5.21; inconspicuousness rank: *t =* 4.03, Tables S3 and S4), most likely because they became accustomed to the set-up and what they were searching for.

The inconspicuousness rank of butterfly species was affected by time of day and the day of the experiment (Fig 1b, Table S5). Species rank was highest earlier in the day (*t =* -1.77). Moreover, older participants omitted fewer butterflies at later hours (*t =* 1.32).

Participants were more likely to find opaque butterflies than transparent ones, following the order *H. ninonia* (H), *C. tutia* (C), *B. seba* (B) and *I. salapia* (I) (H>C, I, B: number of butterflies: *z =* 5.73; inconspicuousness rank: *t =* -7.11; C>B: number of butterflies: *z =* 0.03; inconspicuousness rank: *t =* -1.65; B>I: number of butterflies: *z =* 2.37, Table S4; inconspicuousness rank: *t =* 2.68, Fig 1b). However, the gain in detection with increasing time spent searching was highest for the most transparent species (*z* = -2.75, Fig 2, Fig S7). *H. ninonia* was also the species most frequently found first, followed by *C. tutia, B. seba* and *I. salapia* (*X*^2^ = 19.5, df = 3, *p* < 0.001, Table S3).

**Figure 2.**
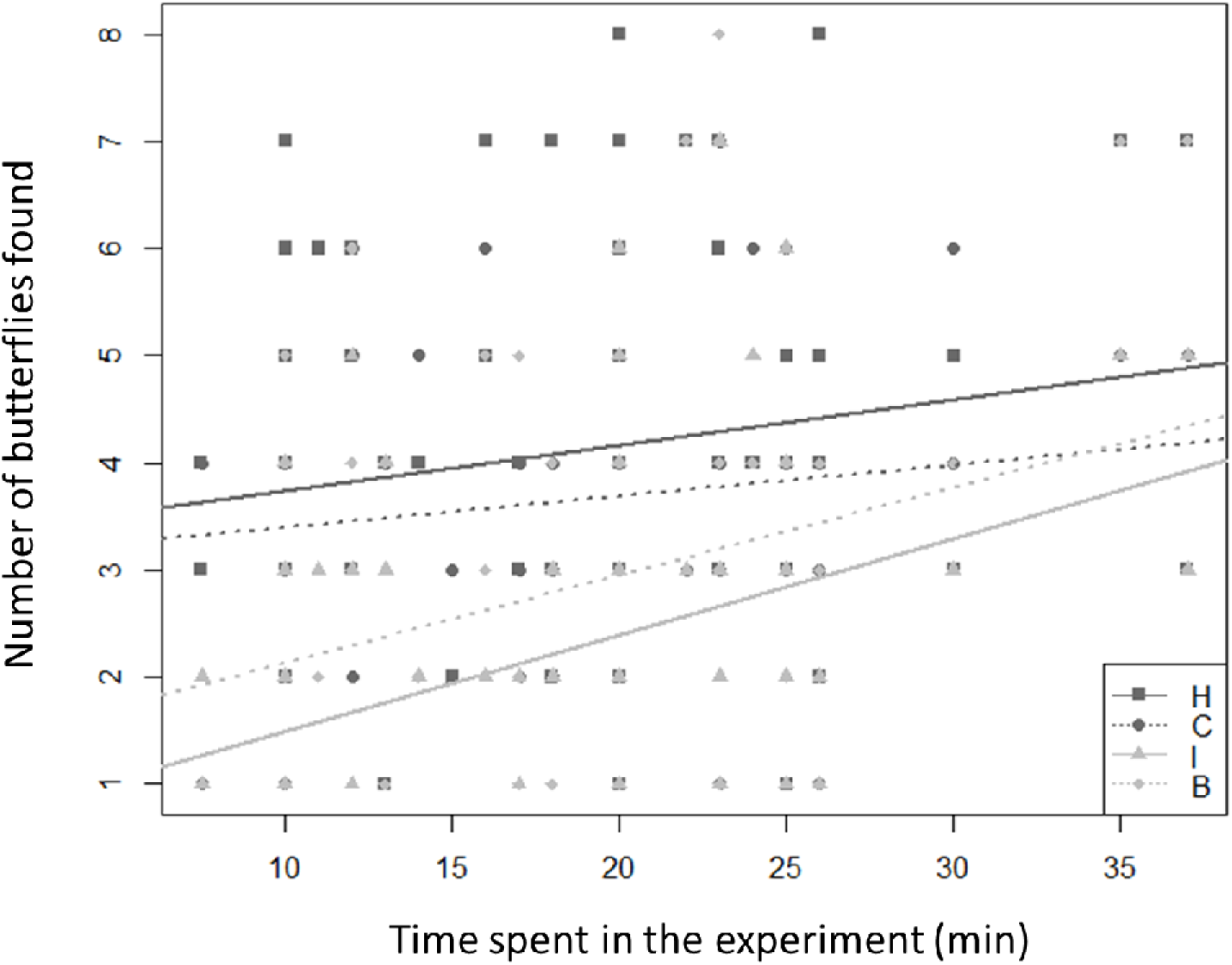
Number of butterflies found for each species according to the time spent completing the experiment by human participants. As shown by the regression lines, spending more time on the experiment resulted in higher numbers of butterflies found, especially for the transparent species (H: estimate slope= 0.043, r^2^ = 0.014, *p* = 0.12; C: estimate slope= 0.03, r^2^ = 0.005, *p* = 0.22; I: estimate slope= 0.090, r^2^ = 0.12, *p* < 0.001; B: estimate slope= 0.08, r^2^ = 0.07, *p* = 0.003). Letters in the legend stand for species names: *H.ninonia* (H), *C.tutia* (C), *I. salapia* (I), and *B. seba* (B). Butterfly transparency increases from top to bottom of the legend (i.e. H<C<I<B).

Differences were also found in the consecutive order in which butterflies were found. Participants were more likely to find two consecutive butterflies of the same species when they were colourful (*H. ninonia* -50 times- and *C. tutia* -58 times) than when they were transparent (*B. seba* -32 times- or *I. salapia* -18 times; *X*^2^ = 29.14, df = 3, *p* < 0.001). *B. seba* and *H. ninonia* were found up to four consecutive times in a single trial. Some species were also more likely to be found consecutively after another species. The most opaque butterflies *H. ninonia* and *C. tutia* (found 278 times consecutively), and the transparent species *B. seba* and *I. salapia* (found 186 times consecutively), were found consecutively more frequently than any of the other possible combinations after correcting for the number of butterflies found for each species (*X*^2^ = 170.95, df = 5, *p* < 0.001). These observed frequencies significantly differed from the position that butterflies occupied on the original set-up along the path (*X*^2^ = 79.12, df = 11, *p* < 0.001, Fig S6b).

### Models of bird and human vision

The achromatic weighted contrast between butterfly colour patches and green-leaf background were similar for both birds and humans (mean achromatic contrast for birds: H=3.81, C= 3.15, I=2.31, B=2.11; for humans: H=5.25, C=4.35, I=3.58, B=3.86. Fig S8). For both observers, *H. ninonia* followed by *C. tutia* (colourful and transparent butterfly) contrasted the most against the leaves, while transparent butterflies (*I. salapia* for humans and *B. seba* for birds), were the least contrasting. Butterflies seem to be more chromatically detectable by birds than for humans (mean chromatic contrast for humans: H = 0.44, C = 0.37, I = 0.25, B = 0.22). For the chromatic contrast seen by birds, *C. tutia,* and then *H. ninona* was the most contrasting, whereas *B. seba* and finally *I. salapia* were the least (mean chromatic contrast for birds: H = 2.02, C = 2.05, I = 1.30, B = 1.38).

## Discussion

### Transparency reduces detectability

As initially predicted based on wing transmittance, and as demonstrated by our behavioural experiments and visual modelling results, transparency decreases butterfly detectability. Interestingly, detection by human participants was similar to naïve birds, as has been shown in other studies (Beatty, Bain, & Sherratt, 2005; Sherratt, Whissell, Webster, & Kikuchi, 2015), providing further support to using human participants to measure predator detection. Surprisingly, experimental results from the bird experiments differed slightly from predictions made based on the measures of transmittance of transparent patches and results obtained from the vision models. For instance, according to the transmittance and the chromatic contrast measured between butterflies and their background, birds should have probably detect *C. tutia* more easily than the two more transparent species. Indeed, semi-transparent objects should be more easily detected than fully transparent objects at short distances and when more light is available (Johnsen & Widder, 1998), such as experimental conditions present during bird experiments. Yet this transparent but brightly coloured species was detected at similar rates as the most transparent species. One possible explanation is that this species possesses disruptive colouration; indeed, wing contours of this species are less strongly delimited than that of the other species and a disrupted outline may hamper detection (Honma, Mappes, & Valkonen, 2015; Stevens & Cuthill, 2006). These contradicting results highlight the importance of combining both modelling and behavioural experiments to better understand the evolution of transparency and other prey defences.

### Transparency in a toxic butterfly?

Our results demonstrate that transparency can effectively reduce prey detectability in chemically-protected ithomiine butterflies. This is surprising as aposematic colour patterns, rather than inconspicuousness, are more common in toxic and unpalatable prey (Mappes et al., 2005; Poulton, 1890; Ruxton, Sherratt, & Speed, 2004). In fact, conspicuousness is often positively correlated with toxicity or unpalatability and can thus be an honest indicator of prey defences (Arenas, Walter, & Stevens, 2015; Blount, Speed, Ruxton, & Stephens, 2009; Maan & Cummings, 2012; Prudic, Skemp, & Papaj, 2007; Sherratt & Beatty, 2003), and predators learn faster to avoid unpalatable prey when colours are more conspicuous (Gittleman & Harvey, 1980; Lindstrom, Alatalo, Mappes, Riipi, & Vertainen, 1999). This might suggest that the evolution of transparency in these butterflies is the result of a loss in unpalatability. If this is the case, the existence of mimicry rings of transparent clearwing butterflies remains unexplained, as this is usually the result of convergence of warning signals promoted by the positive frequency-dependent selection exerted by predators (Willmott et al., 2017). Alternatively, if defences are costly, prey may invest in either visual or chemical defences (Darst, Cummings, & Cannatella, 2006; Speed & Ruxton, 2007; Wang, 2011). Such strategies have been shown to afford equivalent avoidance by predators (Darst et al., 2006). Transparency may therefore be associated with an increase in unpalatability. Unfortunately, the relationship between transparency and the degree of chemical defences in clearwing butterflies is yet unknown.

Alternatively, transparency may lower detection and function as a primary defence, with aposematism taking over as a secondary defence if the prey is detected. Indeed, transparent butterflies were not completely cryptic for either birds or humans. In fact, birds found a similar number of both colourful and transparent butterflies, and humans appear to learn to detect and perhaps remember common elements between the more transparent species, what might be the result of a search image. As such, Ithomiini butterflies may be cryptic from afar, but perceived as conspicuous from up close (Gamberale-Stille, Bragee, & Tullberg, 2009; Tullberg, Merilaita, & Wiklund, 2005). A dual strategy of crypsis and conspicuousness has been described for other prey, including defended prey (Järvi, Sillén-Tullberg, & Wiklund, 1981; Kang, Zahiri, & Sherratt, 2017; Sillén-Tullberg, 1985) For example, toxic salamanders of the genus *Taricha* are generally cryptic, only revealing their warning coloured underbelly when threatened (Johnson & Brodie Jr, 1975). In Ithomiini, conspicuous (or potentially conspicuous) elements can be found on even transparent species, and most species possess opaque areas delineating the edges and contrasting with the background, most likely increasing detection (Stevens & Cuthill, 2006). This, combined with our results and the occurrence of co-mimics in natural populations, suggests that these butterflies may reduce the cost of conspicuousness using transparency in addition to maintaining the benefits of detectable warning signals. Behavioural experiments testing the distance at which Ithomiini butterflies are detected are needed to shed further light on the function of aposematism in less conspicuous prey.

Finally, transparency may have evolved as an additional primary protection against birds such as adult kingbirds (*Tyrannus melancholicus,* C. E. G. Pinheiro, 1996) which are able to tolerate their chemical defences. Indeed, both models (Endler & Mappes, 2004) and experiments (Mappes, Kokko, Ojala, & Lindström, 2014; Valkonen et al., 2012) have shown that weak warning signals (not overtly conspicuous) can evolve and be maintained in communities where predators vary in their probability of attacking defended prey. Larvae of *Dryas julia* butterflies, pine sawfly larvae (*Neodiprion sertifer* for example), and shield bugs (Acanthosomatidae, Heteroptera) are some of the several examples of unpalatable species that display weak visual warning signals (see Endler & Mappes, 2004). Similar to the polymorphic poison frog *Oophaga granulifera*, clearwing species may reflect a continuum between aposematic and cryptic strategies, possibly shaped by differences in the strength of predator selection as a result of the frequency of naïve predators and/or the variation in predator sensitivities to chemical compounds (Willink, BrenesLJMora, Bolaños, & Pröhl, 2013). A thorough characterization of unpalatability, microhabitat and predator communities would be useful in better understanding conditions that promote the evolution of transparency in Ithomiini.

## Conclusions

Our study, combining behavioural experiments with different predators and vision modelling, provides comparative insights into the complex role transparency may play in anti-predator defences of aposematic organisms. We show for the first time that transparency is an effective strategy for the reduction of detectability of terrestrial prey. We also demonstrate that Ithomiini butterflies may in fact be decreasing the costs of conspicuousness, while still retaining visual elements that are recognised as warning signals. Future studies exploring the efficiency of combining transparency and warning signals in decreasing predation risk will further contribute to understanding the evolution of cryptic elements in aposematic prey.

## Acknowledgments

We thank Tuuli Salmi and Tiffanie Kortenhoff for their invaluable help with behavioural experiments, Helinä Nisu for her advice on bird care, SERFOR, Proyecto Huallaga and Gerardo Lamas for providing research permits in Peru (collecting and exportation permit 002-2015-SERFOR-DGGSPFFS), as well as Corentin Clerc, Monica Monllor, Alexandre Toporov and Marc Toporov-Elias for help with collecting butterflies used in this study, Céline Houssin for Ithomiini pictures and calculation of wing surfaces for each patch, Konnevesi Research Station which provided the facilities used for bird experiments, and visitors to Montpellier botanical garden for their enthusiastic contribution. The Academy of Finland to JM, SG and ON (Grants 2100000256 and 21000038821), the Clearwing ANR program (ANR-16-CE02-0012) and the Human Frontier Science Program grant (RGP 0014/2016) fund the study.

## Author contribution

DG, ME, JM and MA designed the study; ME, MM and DG collected the butterfly samples; MA, SG, ON, ME and JM did the experiments; DG and CD took the optical measurements; MA, DG and ME analysed the data; MA, DG, MM, ME, ON, SG and JM wrote the manuscript.

## References

Anderson, D., Burnham, K., & White, G. (1998). Comparison of Akaike information criterion and consistent Akaike information criterion for model selection and statistical inference from capture-recapture studies. Journal of Applied Statistics, 25(2), 263–282.

Arenas, L. M., Walter, D., & Stevens, M. (2015). Signal honesty and predation risk among a closely related group of aposematic species. Scientific Reports, 5.

Beatty, C. D., Bain, R. S., & Sherratt, T. N. (2005). The evolution of aggregation in profitable and unprofitable prey. Animal Behaviour, 70(1), 199–208.

Beccaloni, G. (1997). Ecology, natural history and behaviour of the Ithomiinae Butterflies and their mimics in Ecuador. Tropical Lepidoptera, 8, 103–124.

Blount, J. D., Speed, M. P., Ruxton, G. D., & Stephens, P. A. (2009). Warning displays may function as honest signals of toxicity. Proceedings of the Royal Society of London B: Biological Sciences, 276(1658), 871–877. doi:10.1098/rspb.2008.1407

Brower, L. P., Brower, J. V. Z., & Collins, C. T. (1963). Experimental studies of mimicry: Relative palatability and Müllerian mimicry among Neotropical butterflies of the subfamily Heliconiinae. New York Zoological Society.

Brown, K. S. J. (1984). Chemical ecology of dehydropyrrolizidine alkaloids in adult Ithominae(Lepidoptera: Nymphalidae). Revista Brasileira de Biologia, 44(4), 435–460.

Brown, K. S. J. (1985). Chemical ecology of dehydropyrrolizidine alkaloids in adult Ithomiinae (Lepidoptera: Nymphalidae). Rev. Bras. Biol., 44, 453–460.

Brown, K. S. J., & Neto, J. V. (1976). Predation on aposematic ithomiine butterflies by tanagers (Pipraeidea melanonota). Biotropica, 136–141.

Darst, C. R., Cummings, M. E., & Cannatella, D. C. (2006). A mechanism for diversity in warning signals: conspicuousness versus toxicity in poison frogs. Proceedings of the National Academy of Sciences, 103(15), 5852–5857.

Davis, A. K., Chi, J., Bradley, C., & Altizer, S. (2012). The redder the better: wing color predicts flight performance in monarch butterflies. PloS One, 7(7), e41323.

Elias, M., Gompert, Z., Jiggins, C., & Willmott, K. (2008). Mutualistic interactions drive ecological niche convergence in a diverse butterfly community. PLoS Biology, 6(12), e300.

Endler, J. A. (1988). Frequency-dependent predation, crypsis and aposematic coloration. Phil. Trans. R. Soc. Lond. B, 319(1196), 505–523.

Endler, J. A., & Mappes, J. (2004). Predator mixes and the conspicuousness of aposematic signals. The American Naturalist, 163(4), 532–547.

Freitas, A. V. L., & Brown, K. S. (2004). Phylogeny of the Nymphalidae (Lepidoptera). Systematic Biology, 53(3), 363–383.

Gamberale-Stille, G., Bragee, C., & Tullberg, B. S. (2009). Higher survival of aposematic prey in close encounters with predators: an experimental study of detection distance. Animal Behaviour, 78(1), 111–116.

Gittleman, J. L., & Harvey, P. H. (1980). Why are distasteful prey not cryptic? Nature, 286(5769), 149–150. doi:10.1038/286149a0

Honma, A., Mappes, J., & Valkonen, J. K. (2015). Warning coloration can be disruptive: aposematic marginal wing patterning in the wood tiger moth. Ecology and Evolution, 5(21), 4863–4874.

Ihalainen, E., Rowland, H. M., Speed, M. P., Ruxton, G. D., & Mappes, J. (2012). Prey community structure affects how predators select for Mullerian mimicry. Proceedings of the Royal Society B-Biological Sciences, 279(1736), 2099–2105. doi:10.1098/rspb.2011.2360

Järvi, T., Sillén-Tullberg, B., & Wiklund, C. (1981). The cost of being aposematic. An experimental study of predation on larvae of Papilio machaon by the great tit Parus major. Oikos, 267–272.

Jiggins C. D., Estrada C., & Rodrigues A. (2004). Mimicry and the evolution of premating isolation in Heliconius melpomene Linnaeus. Journal of Evolutionary Biology, 17(3), 680–691. doi:10.1111/j.1420-9101.2004.00675.x

Jiggins, C. D., Mallarino, R., Willmott, K. R., & Bermingham, E. (2006). The phylogenetic pattern of speciation and wing pattern change in neotropical Ithomia butterflies (Lepidoptera: Nymphalidae). Evolution, 60(7), 1454–1466.

Johnsen, S. (2014). Hide and seek in the open sea: pelagic camouflage and visual countermeasures. Annual Review of Marine Science, 6, 369–392.

Johnsen, S., & Widder, E. A. (1998). Transparency and visibility of gelatinous zooplankton from the northwestern Atlantic and Gulf of Mexico. The Biological Bulletin, 195(3), 337–348.

Johnson, J. A., & Brodie Jr, E. D. (1975). The selective advantage of the defensive posture of the newt, Taricha granulosa. American Midland Naturalist, 139–148.

Kang, C., Zahiri, R., & Sherratt, T. N. (2017). Body size affects the evolution of hidden colour signals in moths. Proc. R. Soc. B, 284(1861), 20171287.

Kerfoot, W. C. (1982). A question of taste: crypsis and warning coloration in freshwater zooplankton communities. Ecology, 63(2), 538–554.

Langsdale, J. (1993). Developmental changes in the opacity of larval herring, Clupea harengus, and their implications for vulnerability to predation. Journal of the Marine Biological Association of the United Kingdom, 73(1), 225–232.

Lindstrom, L., Alatalo, R. V., Mappes, J., Riipi, M., & Vertainen, L. (1999). Can aposematic signals evolve by gradual change? Nature, 397(6716), 249–251. doi:10.1038/16692

Maan, M. E., & Cummings, M. E. (2012). Poison frog colors are honest signals of toxicity, particularly for bird predators. The American Naturalist, 179(1), E1–E14. doi:10.1086/663197

Mallet, J., & Singer, M. C. (1987). Individual selection, kin selection, and the shifting balance in the evolution of warning colours: the evidence from butterflies. Biological Journal of the Linnean Society, 32(4), 337–350.

Mappes, J., Kokko, H., Ojala, K., & Lindström, L. (2014). Seasonal changes in predator community switch the direction of selection for prey defences. Nature Communications, 5.

Mappes, J., Marples, N., & Endler, J. A. (2005). The complex business of survival by aposematism. Trends in Ecology & Evolution, 20(11), 598–603. doi:10.1016/j.tree.2005.07.011

Miaoulis, I. N., & Heilman, B. D. (1998). Butterfly thin films serve as solar collectors. Annals of the Entomological Society of America, 91(1), 122–127.

Ödeen, A., Håstad, O., & Alström, P. (2011). Evolution of ultraviolet vision in the largest avian radiation - the passerines. BMC Evolutionary Biology, 11(1), 313. doi:10.1186/1471-2148-11-313

Penney, H. D., Hassall, C., Skevington, J. H., Abbott, K. R., & Sherratt, T. N. (2012). A comparative analysis of the evolution of imperfect mimicry. Nature, 483(7390), 461–464.

Pinheiro, C. E. G. (1996). Palatability and escaping ability in neotropical butterflies: Tests with wild kingbirds (Tyrannus melancholicus, Tyrannidae). Biological Journal of the Linnean Society, 59(4), 351–365. doi:10.1111/j.1095-8312.1996.tb01471.x

Pinheiro, C. E., Medri, Í. M., & Salcedo, A. K. M. (2008). Why do the ithomiines (Lepidoptera, Nymphalidae) aggregate? Notes on a butterfly pocket in central Brazil. Revista Brasileira de Entomologia, 52(4), 610–614.

Poulton, E. B. (1890). The colours of animals: their meaning and use, especially considered in the case of insects. D. Appleton.

Prudic, K. L., Skemp, A. K., & Papaj, D. R. (2007). Aposematic coloration, luminance contrast, and the benefits of conspicuousness. Behavioral Ecology, 18(1), 41–46. doi:10.1093/beheco/arl046

R Foundation for Statistical Computing, R. C. (2014). R: A language and environment for statistical computing. Vienna, Austria.

Ruxton, G. D., Sherratt, T. N., & Speed, M. P. (2004). Avoiding attack: the evolutionary ecology of crypsis, warning signals and mimicry. Oxford University Press.

Sherratt, T. N., & Beatty, C. D. (2003). The evolution of warning signals as reliable indicators of prey defense. The American Naturalist, 162(4), 377–389.

Sherratt, T. N., Whissell, E., Webster, R., & Kikuchi, D. W. (2015). Hierarchical overshadowing of stimuli and its role in mimicry evolution. Animal Behaviour, 108, 73–79.

Sillén-Tullberg, B. (1985). Higher survival of an aposematic than of a cryptic form of a distasteful bug. Oecologia, 67(3), 411–415. doi:10.1007/BF00384948

Speed, M. P., & Ruxton, G. D. (2007). How bright and how nasty: explaining diversity in warning signal strength. Evolution, 61(3), 623–635.

Stevens, M., & Cuthill, I. C. (2006). Disruptive coloration, crypsis and edge detection in early visual processing. Proceedings of the Royal Society B: Biological Sciences, 273(1598), 2141. doi:10.1098/rspb.2006.3556

Suzuki, T. K., Tomita, S., & Sezutsu, H. (2014). Gradual and contingent evolutionary emergence of leaf mimicry in butterfly wing patterns. BMC Evolutionary Biology, 14(1), 229. doi:10.1186/s12862-014-0229-5

Tsuda, A., Hiroaki, S., & Hirose, T. (1998). Effect of gut content on the vulnerability of copepods to visual predation. Limnology and Oceanography, 43(8), 1944–1947.

Tullberg, B. S., Merilaita, S., & Wiklund, C. (2005). Aposematism and crypsis combined as a result of distance dependence: functional versatility of the colour pattern in the swallowtail butterfly larva. Proceedings of the Royal Society of London B: Biological Sciences, 272(1570), 1315–1321.

Valkonen, J. K., Nokelainen, O., Niskanen, M., Kilpimaa, J., Björklund, M., & Mappes, J. (2012). Variation in predator species abundance can cause variable selection pressure on warning signaling prey. Ecology and Evolution, 2(8), 1971–1976.

Wanasekara, N. D., & Chalivendra, V. B. (2011). Role of surface roughness on wettability and coefficient of restitution in butterfly wings. Soft Matter, 7(2), 373–379.

Wang, I. J. (2011). Inversely related aposematic traits: reduced conspicuousness evolves with increased toxicity in a polymorphic poisonldart frog. Evolution: International Journal of Organic Evolution, 65(6), 1637–1649.

Watson, G. S., Myhra, S., Cribb, B. W., & Watson, J. A. (2008). Putative functions and functional efficiency of ordered cuticular nanoarrays on insect wings. Biophysical Journal, 94(8), 3352–3360.

Willink, B., BreneslMora, E., Bolaños, F., & Pröhl, H. (2013). Not everything is black and white: color and behavioral variation reveal a continuum between cryptic and aposematic strategies in a polymorphic poison frog. Evolution, 67(10), 2783–2794.

Willmott, K. R., Willmott, J. C. R., Elias, M., & Jiggins, C. D. (2017). Maintaining mimicry diversity: optimal warning colour patterns differ among microhabitats in Amazonian clearwing butterflies (Vol. 284, p. 20170744). Presented at the Proc. R. Soc. B, The Royal Society.

Yoshida, A., Motoyama, M., Kosaku, A., & Miyamoto, K. (1997). Antireflective nanoprotuberance array in the transparent wing of a hawkmoth, Cephonodes hylas. Zoological Science, 14(5), 737–741.

Zaret, T. M. (1972). Predators, invisible prey, and the nature of polymorphism in the cladocera (class Crustacea). Limnology and Oceanography, 17(2), 171–184. doi:10.4319/lo.1972.17.2.0171

